## Supplemental Appendix S1 for "Pharmacodynamic model of the dynamic response of *Pseudomonas aeruginosa* biofilms to drug treatments"

### S1 Appendix: Diffusive flux to biofilm cells

The pharmacodynamic model for biofilm cell killing accounts for the dynamics of drug availability at the biofilm cells, which is assumed to be controlled by diffusive flux from the bulk. Note that diffusion can occur both through a hydrodynamic boundary layer, because drug is convected through the flow cell chamber, and through the layer of the biofilm itself. Since the thickness of the biofilm is unknown, these two diffusive regions are lumped together as one.

The flux is determined from the one-dimensional diffusion equation:

$$\frac{\partial c}{\partial t} = \mathcal{D}\frac{\partial^2 c}{\partial x^2}, \tag{A1}$$

where $c$ is the concentration of drug at depth in the boundary layer, $x$, at time $t$, and $D$ is the effective diffusivity of the drug. This partial differential equation is subject to initial and boundary conditions:

$$@t = 0, c = 0 \tag{A2}$$

$$@x = H, c = c_0$$

$$@x = 0, c = 0,$$

where $H$ is the depth of the boundary layer. By scaling the equations according to $\theta = c/c_0$, $\xi = x/H$, and $\tau = Dt/H^2$, equation (A1) becomes:

$$\frac{\partial \theta}{\partial \tau} = \frac{\partial^2 \theta}{\partial \xi^2}, \tag{A3}$$

and the boundary conditions:

$$@\,\tau = 0, \theta = 0$$

$$@\,\xi = 1, \theta = 1 \tag{A4}$$

$$@\,\xi = 0, \theta = 0.$$

This equation and boundary conditions give rise to a Fourier series solution:

$$\theta = \xi + \frac{2}{\pi} \sum_{n=1}^{\infty} \frac{(-1)^n}{n} \sin(n\pi\xi)\, e^{-n^2\pi^2\tau}\,. \tag{A5}$$

The desired flux is found from Fick's Law applied to the cell interface:

$$J = \mathcal{D} \frac{\partial c}{\partial x}\bigg|_{x=0} = \frac{\mathcal{D} c_0}{H} \frac{\partial \theta}{\partial \xi}\bigg|_{\xi=0} = \frac{\mathcal{D} c_0}{H} \left[1 + 2 \sum_{n=1}^{\infty} (-1)^n e^{-n^2\pi^2\tau}\right]. \tag{A6}$$

For experiments in which drug is introduced transiently, the diffusion equation is solved in two time domains. The first, which applies from time 0 to $t^*$, is governed by the above equations and boundary conditions. For the second domain, from time $t^*$ to $t$, the dimensionless time, $\tau'$, is the time since the drug supply was turned off, $\tau' = D\,(t - t^*)/H^2$. The initial condition for this domain is the concentration profile corresponding to the solution (A5) at time $t^*$ ($\tau^* = D\,t^*/H^2$). Since the drug supply has been turned off, there is no additional flux from the bulk fluid into the boundary layer, which sets the boundary condition at the boundary layer-bulk fluid interface. That is,

$$\frac{\partial\theta}{\partial\tau'} = \frac{\partial^2\theta}{\partial\xi^2}, \qquad \text{(A7)}$$

is solved subject to the boundary conditions:

$$@\tau' = 0, \theta = g(\xi)$$

$$@\ \xi = 1, \frac{\partial\theta}{\partial\xi} = 1 \qquad \text{(A8)}$$

$$@\ \xi = 0, \theta = 0.$$

The solution to (A7) subject to (A8) is also given by a series solution:

$$\theta = \sum_{m=1,3,5,...}^{\infty} D_m \sin\left(\frac{m\pi\xi}{2}\right) e^{-m^2\pi^2\tau'/4}\ , \qquad \text{(A9)}$$

in which the series coefficients, $D_m$, are determined from the concentration profile at the time the drug supply was turned off:

$$D_m = \frac{2}{\pi^2}\left\{\frac{4}{m^2}\sin\left(\frac{m\pi}{2}\right) + \sum_{n=1}^{\infty}\frac{(-1)^n}{n}e^{-n^2\pi^2\tau'}\left[\frac{\sin[(n-m/2)\pi]}{n-m/2} - \frac{\sin[(n+m/2)\pi]}{n+m/2}\right]\right\} \qquad \text{(A10)}$$

The desired flux is again found from Fick's Law applied to the cell interface:

$$J = \mathcal{D}\frac{\partial c}{\partial x}\bigg|_{x=0} = \frac{\mathcal{D}c_0}{H}\frac{\partial\theta}{\partial\xi}\bigg|_{\xi=0} = \frac{\pi}{2}\sum_{m=1,3,5,...}^{\infty} mD_m e^{-m^2\pi^2\tau'/4}\ . \qquad \text{(A11)}$$
