## Supplemental Table S1 for "Pharmacodynamic model of the dynamic response of *Pseudomonas aeruginosa* biofilms to drug treatments"

**S1 Table: Effect of transit compartment number on error**

**S1 Table 1: Tobramycin**

| | **0 Comp.** | **1 Comp.** | **2 Comp.** | **3 Comp.** | **4 Comp.** | **5 Comp.** | **6 Comp.** |
|---|---|---|---|---|---|---|---|
| $\mu_B$ | 0.0000 | 0.1849 | 0.0185 | 0.0432 | 0.0713 | 0.0321 | 0.0300 |
| $\alpha$ | 0.0351 | 0.0046 | 0.0057 | 0.0005 | 0.0003 | 0.0002 | 0.0005 |
| $\beta$ | 0.02 | 0.1168 | 0.1472 | 0.1021 | 0.1832 | 0.2088 | 0.5713 |
| $\gamma$ | 3.4315 | 1.8425 | 1.6725 | 2.4132 | 3.2428 | 3.5330 | 2.7133 |
| $k_t$ | 0.00 | 0.1481 | 0.3747 | 0.3895 | 0.4495 | 0.5424 | 0.6120 |
| **Error** | 0.6449 | 0.2714 | 0.3469 | 0.0604 | 0.0563 | 0.0561 | 0.0718 |

**S1 Table 2: Colistin**

| | 0 Comp. | 1 Comp. | 2 Comp. | 3 Comp. | 4 Comp. | 5 Comp. |
|---|---|---|---|---|---|---|
| $\mu_B$ | 0.0466 | 0.0001 | 0.062 | 0.1424 | 0.290 | 0.4631 |
| $\alpha$ | 0.0238 | 0.0082 | 0.0191 | 0.023 | 0.0224 | 0.0211 |
| $\beta$ | 0.4165 | 0.3986 | 0.6889 | 1.0170 | 1.765 | 3.3533 |
| $\gamma$ | 2.9409 | 4.4313 | 2.93 | 2.50 | 2.357 | 2.2343 |
| $k_t$ | 0.0000 | 1.8924 | 4.1184 | 4.9999 | 4.9999 | 4.9999 |
| **Error** | 1.4165 | 1.1361 | 1.6946 | 2.0074 | 2.6296 | 3.3053 |
